# Gradient-mixing LEGO robots for purifying DNA origami nanostructures of multiple components by rate-zonal centrifugation

**DOI:** 10.1101/2021.07.02.450731

**Authors:** Jason Sentosa, Brian Horne, Franky Djutanta, Dominic Showkeir, Robert Rezvani, Rizal F. Hariadi

## Abstract

DNA origami purification is critical in emerging applications of functionalized DNA nanostructures from basic fundamental biophysics, nanorobots to therapeutics. Advances in DNA origami purification have led to the establishment of rate-zonal centrifugation (RZC) as a scalable, high-yield, and contamination-free approach to purifying DNA origami nanostructures. In RZC purification, a linear density gradient is created using viscous agents, such as glycerol and sucrose, to separate molecules based on their mass and shape during high-rpm centrifugation. However, current methods for creating density gradients are typically time-consuming because of their reliance on slow passive diffusion. Here, we built a LEGO gradient mixer to rapidly create a quasi-continuous density gradient with minimal layering of concentrations using simple rotational motion. We found that rotating two layers of different concentrations at an angle can reduce the diffusion time from a few hours to mere minutes. The instrument needed to perform the movement can be constructed from low-cost components, such as Arduino and LEGO Mindstorms pieces, and has comparable efficacy to commercial gradient mixers currently available. Our results demonstrate that the creation of a linear density gradient can be achieved with minimal labor, time, and cost with this machine. With the recent advances in DNA origami production, we anticipate our findings to further improve the viability of scaling up DNA origami purification in grams quantities. Our simple process enables automated large-scale purification of functionalized DNA origami more feasible in resource-constrained settings.

## Introduction

**D**NA origami, a facile method to self-assemble complex nanostructures from long, single-stranded DNA (scaffold strands) and programmed shorter oligonucleotides (staple strands),^1–8^ has proven to be a reliable and efficient method to create nanostructures with well-defined shape and customizable geometry. The ability to fabricate the desired shapes and features of a nanoscale object offers a powerful tool for nanotechnology. ^8–11^ The highly specific structure of DNA origami has led to its use in fields such as medicine, ^12–16^ microscopy,^17–20^ and electronics.^21–23^ Although efforts have been made to optimize DNA origami assembly,^24,25^ the yield of well-folded origami is far less than 100%.^26^ Most applications of DNA origami require uncontaminated structures that are free of staple strands and misfolded origami.^26–28^ Hence, a purification step is added to isolate the well-folded structures.

Rate-Zonal Centrifugation (RZC) is a high-yield, contamination-free method of purifying DNA origami.^27^ The technique employs a linear density gradient that is subjected to high centrifugal force to separate heterogeneous molecules based on their distinct mass and shape.^29^ In the context of DNA origami, RZC can be used to separate well-folded structures from other undesired species (misfolded structures, staple strands, and aggregates). RZC has several advantages compared to other purification methods, such as agarose gel electrophoresis (AGE). RZC keeps the samples in aqueous solution throughout the process, is free from contaminants such as agarose gel residues, and is scalable to accommodate a large amount of sample.^27^

Although RZC may take only 1 to 3 hours and minimal labor to perform, preparing the density gradient can be time-consuming.^27,30^ There are several ways to prepare a glycerol density gradient: layering two solutions of different glycerol concentrations in a tube and resting the tube horizontally for 1 to 2 hours to let the glycerol to passively diffuse, overlaying or underlaying several layers of different glycerol concentration and relying on passive diffusion, or mixing the glycerol gradient using a gradient mixer instrument. The first two methods are mostly cost-free, but time-consuming to prepare. Meanwhile, the third option is fast but requires practice and can be costly. Therefore, there is a compelling need to establish a density gradient creation method that is both fast and cost-effective to create.

Here, we report a fast and low-cost method to create a linear glycerol density gradient relying on simple rotational motion to accelerate diffusion. The instrument was made using low-cost materials from LEGO (EV3 Mindstorms). The diffusion process facilitated by the LEGO gradient mixer takes one minute, down from the several-hour process of manual methods. The general process is as follows: A low-concentration solution of glycerol is gently layered on top of a high-concentration glycerol solution inside a centrifuge tube. The glycerol-filled tubes are then loaded into the LEGO gradient mixer and the program is initiated to tilt the glycerol-filled tubes horizontally and rotate them for 1 minute at low rpm (20 rpm). The LEGO gradient mixer will return to its upright position and DNA origami samples (6-hb monomer and dimer) are loaded on top of the newly-created glycerol gradients. The tubes are then transferred to an ultracentrifuge for RZC separation.^27^ Fractions are collected from the RZC result to be analyzed using AGE and the purified structures are verified using atomic force microscopy (AFM).

## Materials and Methods

### Buffers, reagents, materials and equipments

All buffers, reagents, materials, and equipment can be found in Section S4 of the Supporting Information.

### Design of the Gradient Mixer

All components for building the LEGO gradient mixer are part of the EV3 LEGO Mindstorms kit (Item no. 31313) except the ultracentrifuge tube holder (Fig. 1a), which was 3D printed using an Ultimaker 3 3D printer (part no. 9671) (see Supplementary Fig. S1 for building instruction). The design features two motors: A spinning motor (Fig. 1b) and turning servo motor (Fig. 1c). The centrifuge tube holder is connected to the spinning motor, which is connected to the servo motor by the large grey gear (Fig. 1d). Both motors are supported by the scaffold (Fig. 1e), which connects directly to the large grey gear, allowing the spinning motor and the centrifuge tube holder to rotate into a horizontal position. Both motors are connected to the LEGO Brick (Fig. 1f), which was programmed with the protocol. When the LEGO gradient mixer is ready to be used, the servo motor slowly tilts the tubes into a horizontal position. The tubes are tilted horizontally to create greater surface area between the glycerol solutions, which accelerates diffusion. After reaching a horizontal configuration, the spinning motor begins to rotate the tubes. This allows layers of different concentrations to diffuse quickly with a smooth transition. Once the spinning motor stops, the servo motor returns the tubes back to their initial position and the gradient is ready for RZC.

**Fig. 1.**
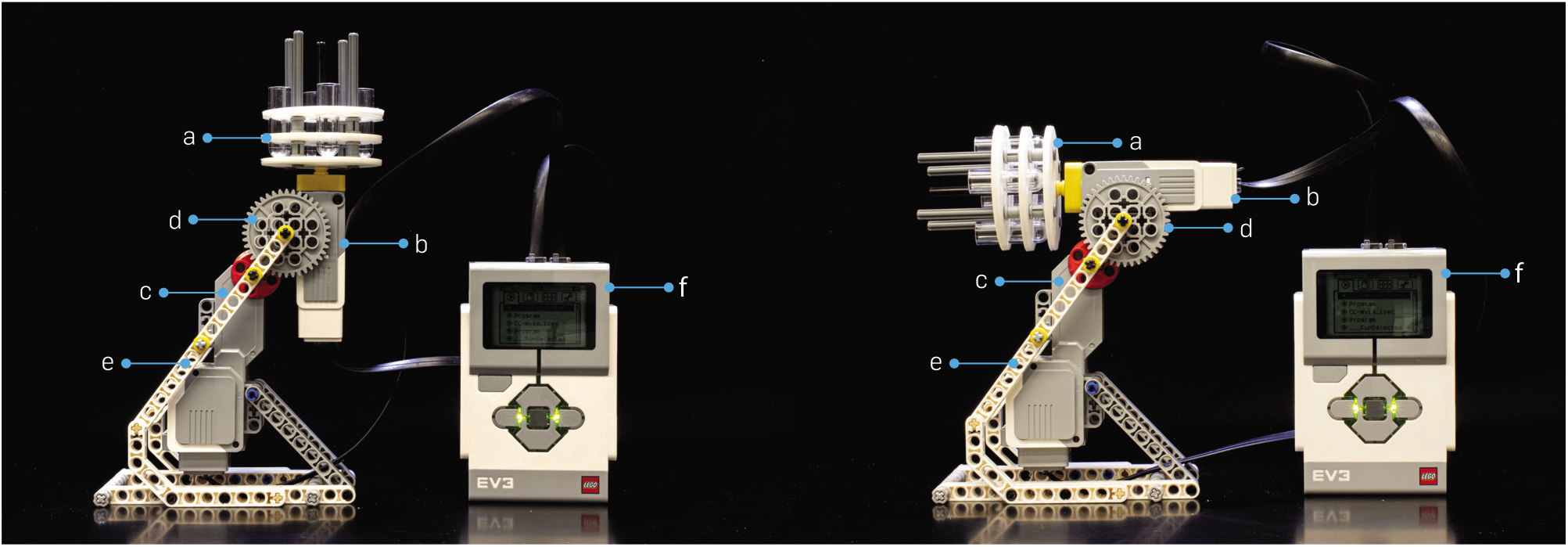
Side view of the LEGO gradient mixer during its initial position (**left**) and its horizontal tilting phase (**right**). (**a**) 3D printed centrifuge-tube holder. (**b**) Spinning motor to rotate the tubes in its horizontal position. (**c**) Turning servo motor responsible for tilting the tubes horizontally. (**d**) Large grey gear connecting the two motors with its small gear complement. (**e**) The scaffold holding the structure together. (**f**) The LEGO controller for orchestrating the motions of the two motors.

### DNA Origami Design and Assembly

The design for monomers and dimers was adopted from Shawn Douglas’s paper,^31^ which utilized a similar structure. The monomer was constructed as a single DNA origami while the dimer was constructed from two separate pieces of precursor monomer (left and right). The left and right precursor monomers were annealed individually with different staple strands such that one side of each precursor monomer was on and the other side was off. The resulting left and right monomers were filtered using a 100 kD amicon filter (4,500 ×g for 5 mins twice) to get rid of excess staples that may have prevented dimerization. The filtered products were then combined into a single tube to dimerize overnight. The CADnano file for both 6-hb monomers and dimers can be found in the Supplementary Materials Repository.

The assembly of the DNA nanostructures was accomplished in a one-pot reaction by mixing 30 nm p8094 scaffold strands and 300–900 nm staple oligonucleotides (Table S1) in a buffer solution containing 1× TAE 12.5 mM MgCl_2_. The mixture was then annealed using a thermocycler that gradually cooled the mixture from 90 °C to 30 °C over 1.5 hours. The predicted length of a 6-hb monomer is 460 nm for 32 segments of 42 bp each, assuming the length of each base pair is 0.34 nm. The dimers have a predicted length of 920 nm with 32 segments from both the left and right part and 2/3 segment for the connectivity (total 642/3 segments) of 42 bp each segment. The diameter of the 6-hb origami calculated by assuming an interhelical distance of 3 nm^32^ is approximately 8 nm.

### Preparation of the Glycerol Gradient

A 15–45% (concentrations depend on the origami of interest) quasi-continuous glycerol gradient was prepared in the following way. First, 300 μL of 45% glycerol (v/v) in 1× TAE 12.5 mM MgCl_2_ buffer was pipetted into a centrifuge tube. The same volume of 15% glycerol solution was then carefully layered on top of the 45% glycerol using either a dropper or a syringe to prevent disturbing the surface of the 45% glycerol (Fig. 2A.1). A clear division between the two solutions was visible. The glycerol filled centrifuge tubes were then inserted into the LEGO gradient mixer and the protocol was run to mix the layered glycerol for 1 minute, which was sufficient for the 1.0 mL tubes (Fig. 2A.2–2A.4).

**Fig. 2.**
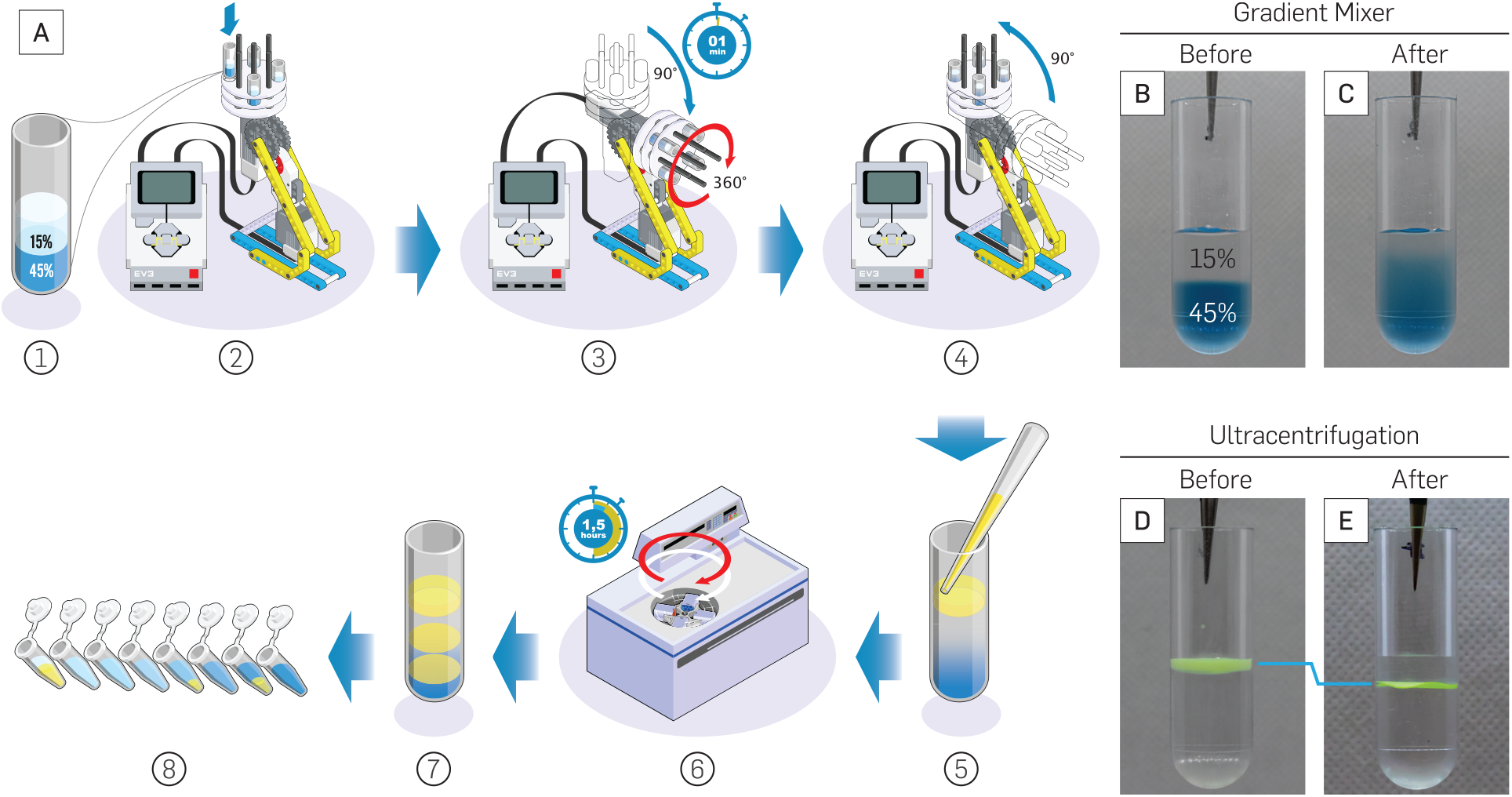
Glycerol gradient preparation and RZC purification (Movie S1). (**A**) Preparation of glycerol gradient with LEGO gradient mixer and separation of sample by RZC. (**B**) Layers of glycerol before mixing; the blue glycerol at the bottom is 45% (v/v) and the clear glycerol at the top is 15% (v/v). (**C**) Linear glycerol gradient after mixing with LEGO gradient mixer. (**D**) 140 nm green fluorescent beads on top of the glycerol gradient before RZC. (**E**) Fluorescent beads after RZC concentrated into a thin layer due to separation from the glycerol gradient. All glycerol solutions were diluted in 1 × TAE 12.5 mM MgCl_2_

The ideal spin time was discovered using a visual test. First, the 45% glycerol solution was dyed with blue food coloring. Both the clear 15% glycerol and the colored 45% glycerol (300 μL each) were added to the centrifuge tubes (Fig. 2B). The tubes were loaded into the mixer and different spin times were tested until a smooth transition from clear (15%) to blue (45%) was visible (Fig. 2C). Three other glycerol gradient concentrations (30%/45%, 15%/60%, 30%/60%) were also tested with varying spin times of 30 sec, 60 sec, 120 sec, and 180 sec. The results showed that one minute of spin time was sufficient to create the desired glycerol gradient. The 30 seconds of spin time produced a step gradient, while 120 seconds and 180 seconds of spin time produced a uniform solution rather than a gradient (Fig. 3). The detailed protocol programmed into the LEGO gradient mixer for both the experiment and the visual test can be found in the Supplementary Materials Repository.

**Fig. 3.**
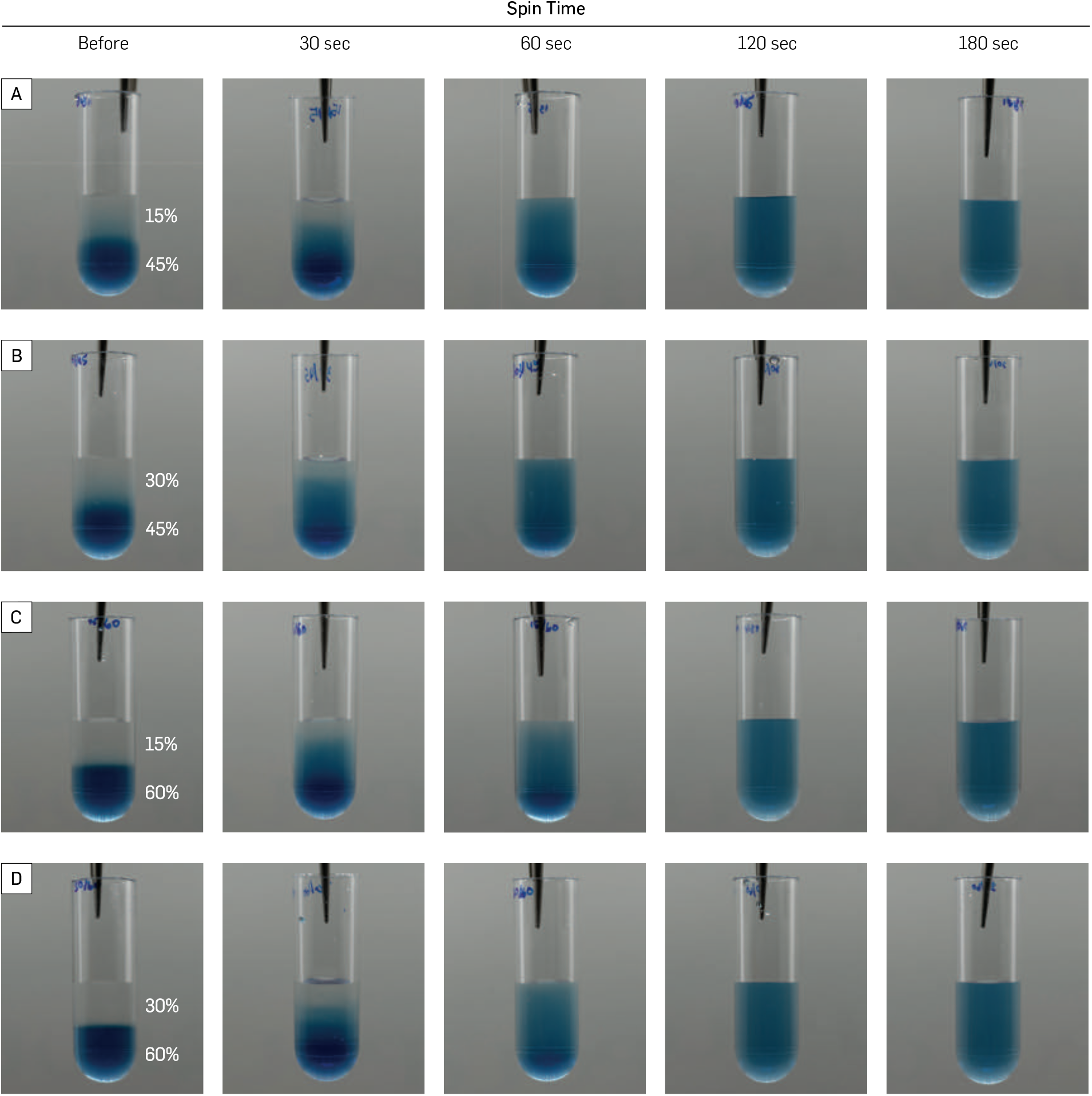
Comparison of different spin times for different gradients. (**A–D**) Four different top and bottom glycerol concentration after 0, 30, 60, 120, and 180 sec. The bottom glycerol is dyed blue for visual inspection, while the top glycerol layer was kept uncolored.

### Centrifugation and Sample Extraction Procedure

After the glycerol gradient was formed, 50-100 μL of the sample containing 10% glycerol was loaded on top of the gradient by hovering the pipette tip near the surface of the gradient (Fig. 2A.5). Most of the sample should rest on the surface of the gradient. The centrifuge tubes containing the sample and gradient were loaded into a swinging bucket rotor (Beckman TLS 55). The samples were centrifuged at 50,000 rpm (~150,000 g) for 1.5 hours at 4° C (Fig. 2A.6). For different origami, the exact RPM and duration of centrifugation need to be optimized. Once the centrifugation was complete, 16 to 18 equal-volume fractions of the sample (Fig. 2A.7–8) were then collected from bottom to top using longneck gel-loading tips to preserve the glycerol layers (see Supplementary Movie S1 for full instructional video from gradient preparation to fractionation). Fluorescent beads (G140) were used as a control sample for RZC protocol. The beads were laid on top of the 15–45% glycerol gradient (Fig. 2D) to be centrifuged with the above protocol. The resulting tube showed the beads concentrating into a thin layer that lies below the surface where it initially was (Fig. 2E).

### Experiment Calibration

Optimization of gradient parameters and centrifugation setup were done using SYBR Gold-stained sample for quick separation assessment. The sample being purified was stained with 3× SYBR Gold prior to loading into the experimented gradients. The centrifuge tubes containing the stained sample and gradient were centrifuged with the experimented setup. After centrifugation, the resulting tubes were visualized with Invitrogen Safe Imager 2.0 Blue Light transilluminators and the images were taken with a Canon lens EF-S 35mm f/2.8 Macro IS STM covered with a single B+W 49mm XS-Pro clear MRC-nano 007 filter on a Canon EOS 77D camera body. The structures being separated showed a thin horizontal band on the gradient while the staple strands showed a cloudy region near the top of the gradient. The best parameters were then determined by looking at the farthest separation distance between the structures being purified and the undesired molecules.

### Result Confirmation and Analysis

After the centrifuged sample gradient had been partitioned into fractions, 10 μL from each fraction was taken and mixed with 1 μL of 20× SYBR gold and 1 μL of loading dye. The fractionated sample was analyzed using AGE. 6-hb monomers underwent gel electrophoresis in 1% Agarose Gel at 75V for 1.5 hours. Once the electrophoresis was complete, the gel was post-stained with 1× SYBR gold and imaged using ultraviolet light in Bio-Rad Molecular Imager Gel Doc XR+ (1708195EDU). The AGE was done to detect the fractions containing the desired origami, which were then combined into a single tube. Buffer exchange on the purified origami is recommended to ensure the structures stay in their native buffer without any of the glycerol.

The result of the gradient purification was further confirmed by AFM imaging to compare purified and unpurified samples. Both purified and unpurified samples were diluted by 100× with 1× TAE 12.5 mM MgCl_2_ to ensure the DNA nanotubes were sparsely distributed in the AFM scans. After preparing and calibrating the AFM, four samples were imaged: unpurified 6-hb monomer, purified 6-hb monomer, unpurified 6-hb dimer, and purified 6-hb dimer.

### Statistical Analysis

The 6-hb monomer and dimer were distinguishable by their length in the AFM images, ~500 nm for a monomer and ~1000 nm for a dimer. The AFM images were processed in imageJ and the number of monomer and dimer were counted to calculate the content of the purified and unpurified dimers. The bootstrap method was used to calculate the standard error of mean of the content (Fig. 5I–J). The bootstrap calculation was performed with either MATLAB or Mathematica. First, the monomer and dimer were assigned as binary number 0 and 1 respectively. Then, from the full data set of sample *N*, a subset of size *N/2* was randomly chosen to compute the mean. The size of the subset was rounded to the nearest integer and an element was never chosen more than once. 10,000 subsets and means were generated with randomly chosen elements each time. Finally, the standard deviation of the means were used to estimate the standard error of the monomer and dimer content.

**Fig. 4.**
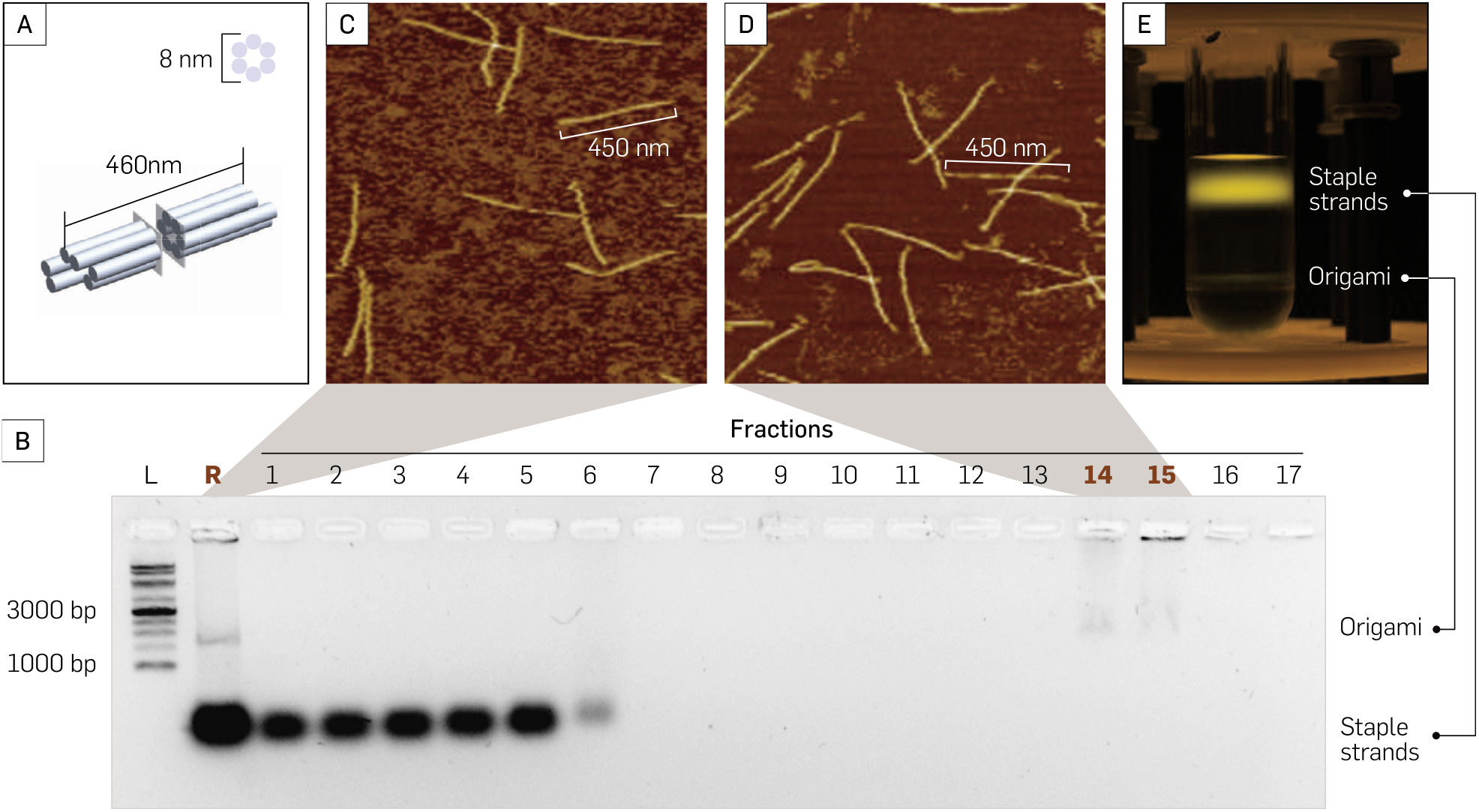
RZC purification result for 6-hb monomer. (**A**) 3D rendered model of 6-hb monomer (not to scale; created using Solidworks). (**B**) Gel result of monomer fractions from top (1) to bottom (17). *L* is 1 kb ladder (Goldbio 1kb ladder), *R* is raw unpurified monomer, underlined fraction numbers indicate the fractions containing the purified monomer. (**C**) AFM image of the unpurified monomer. (**D**) AFM image of the RZC purified monomer. (**E**) SYBR gold stained monomer after being separated with RZC.

**Fig. 5.**
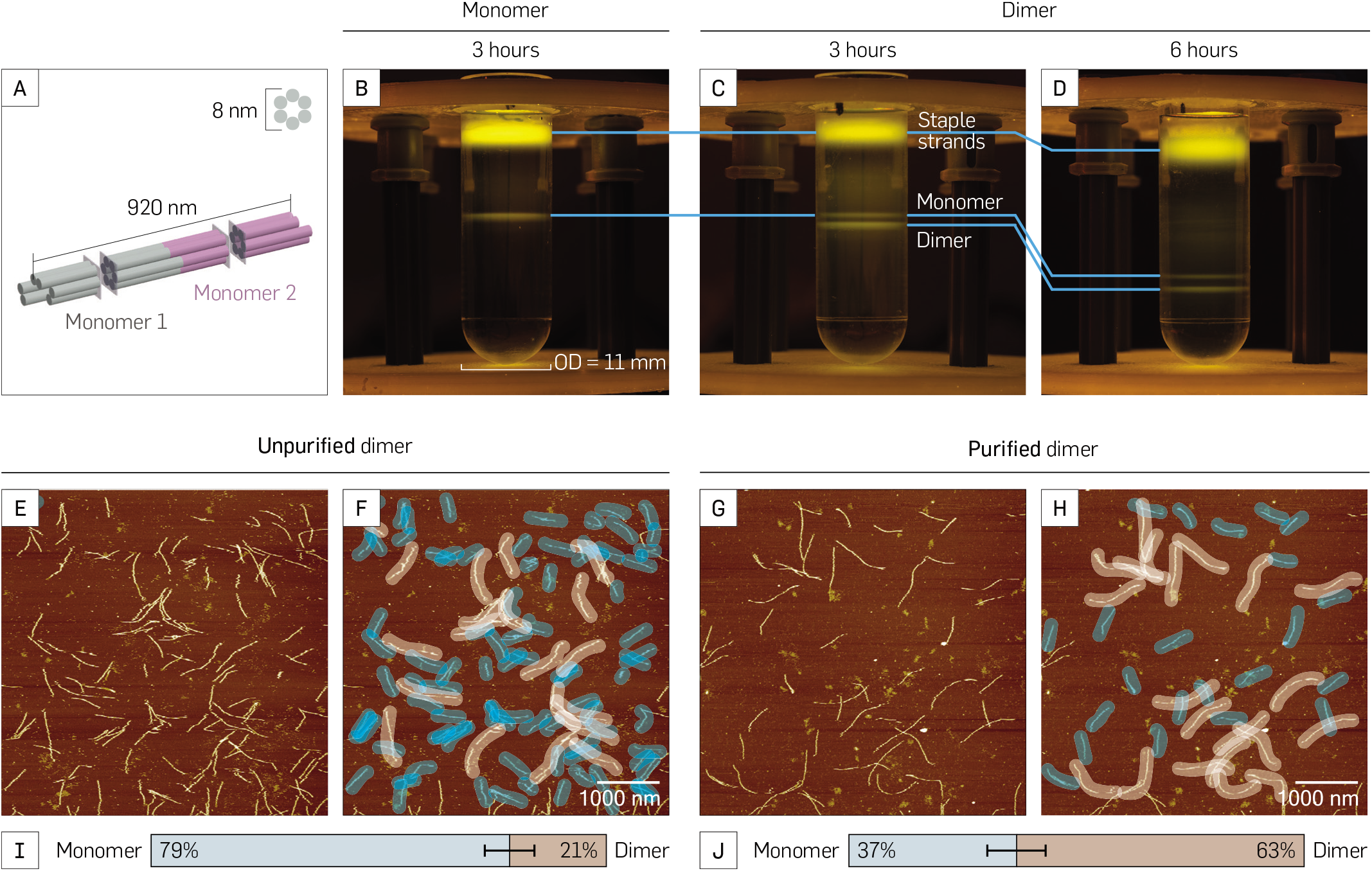
RZC purification of 6-hb dimer. (**A**) 3D-rendered model of a 6-hb dimer (not to scale). (**B**) Comparison between SYBR gold stained 6-hb monomer (left) and (**C**) 6-hb dimer (right) purified using a 30% – 60% gradient column centrifuged for 3 hours at 50,000 rpm in 4 °C. (**D**) SYBR gold stained dimer with 6 hours of centrifugation. (**E**) AFM image of unpurified dimer. (**F**) Precursor monomer (highlighted blue) and dimer (highlighted light brown) from unpurified dimer. (**G**) AFM image of RZC purified dimer. (**H**) Significantly smaller number of monomer (highlighted blue) with similar number of dimer (highlighted light brown) compared to those in unpurified dimer. (**I** and **J**) Comparison of precursor monomers (highlighted blue) and dimer (highlighted light brown) between unpurified and purified dimer. Dimer content of unpurified dimer and purified dimer at 21±5% and 63±6% respectively. Standard deviation was calculated with the bootstrapping method (*N* =273 for unpurified dimer; *N* =114 for purified dimer).

## Results

### Lighter staples remain in the top section after purification

Two DNA origami structures, a 6-hb monomer (Fig. 4A) and 6-hb dimer (Fig. 5A), were purified using the glycerol gradient using the LEGO gradient mixer. The effectiveness of the gradient used for rate-zonal centrifugation was evaluated using AGE analysis and AFM imaging. The sample separation after fractionation between staples, misfolded structures, and well-folded structures was comparable to the results of other glycerol gradient preparation methods.^27^ The AGE results (Fig. 4B) showed that the well-folded structures were contained in 10–20% of the glycerol gradient volume and most of the staples were accumulated near the top of the gradient, considerably separated from the origami of interest.

AFM images of the 6-hb monomer also revealed that most of the staple strands were removed after purification. The unpurified monomer AFM image surface is crowded with specs of staple strands that reside in the mixture (Fig. 4C). Conversely, the purified monomer AFM surface is much cleaner (Fig. 4D).

### SYBR gold staining enables visualization of separated DNA origami

The SYBR gold-stained 6-hb monomer sample clearly showed the separation between the well-folded monomer and its excess staples using a common glycerol gradient concentration of 15% to 45% with 50,000 rpm ultracentrifuge at 4 °C for 1.5 hours (Fig. 4E).

Meanwhile, the 6-hb dimer did not show separation between the dimer and its precursor monomer with the same experimental conditions as the monomer. Fine-tuning of the experiment was done with the help of an SYBR gold visualization test. It was determined that a glycerol gradient concentration of 30% to 60% (600 μL each) spun at 50,000 rpm at 4 °C was able to separate the dimer and its precursor monomer. The dimer was compared to the monomer in the same gradient condition after 3 hours of centrifugation at 50,000 rpm at 4 °C (Fig. 5B–C). The monomer separation yielded only one thin band which corresponded to the 6-hb monomer. On the other hand, the dimer showed two bands: the upper band that was near the same level as the monomer corresponded to the precursor monomer of the dimer; the lower band corresponded to the 6-hb dimer, which has twice the mass and size of the monomer, allowing it to migrate further than the monomer during RZC. The dimer showed more separation distance from the monomer after another 3 hours of centrifugation, a total of 6 hours (Fig. 5D).

### 6-hb monomer structure was preserved after purification

Inspection of the DNA origami using AFM revealed that the monomer structures were not compromised (Fig. 4D) by purification. The 6-hb monomer was neither degraded nor physically altered after the purification step. This verification is important because functional DNA origami is needed for most applications. If the DNA structures are not compatible with glycerol, a buffer exchange is needed for long-term storage of the structures. Alternatively, sucrose can be used to substitute glycerol. A sucrose gradient created using the LEGO gradient mixer was able to separate the dimer with similar effectiveness to the glycerol gradient (see Figs. S2 and S3 for dimer separation using sucrose gradient).

### Purification of 6-hb dimers from monomers

The AFM images reveal that separation of the dimer from its precursor monomer using the gradient from the LEGO gradient mixer was partially successful (see Supplementary Materials Repository for more AFM images of purified and unpurified 6-hb dimer). Comparison between the unpurified and purified dimer AFM images showed that there are more monomer-like structures in the unpurified dimer (79±5%; Fig. 5E–F) than in the purified dimer (37±6%; Fig. 5G–H). Percentage of DNA origami dimer over the sum of all the origami (precursor monomers and dimer) showed that the unpurified dimer (Fig. 5I) have a significantly lower dimer to monomer percentage compared to the purified dimer (Fig. 5J), 21±5% and 63±6% respectively. The non-zero monomer quantity in the dimer fraction (Fig. 5C and D) is likely due to the fluid flow during the extraction of the monomer and dimer fractions using an automatic micropipette with a long neck pipette tip.

## Discussion

DNA nanotechnology has the potential to solve real-world problems in the fields of microscopy,^17–20^ nanomedicine,^12–16^ and even molecular-scale electronics.^21–23^ Its pace toward industry applications is paired with successful efforts to remove the key obstacle of scalability for both production^33–35^ and purification^27^ of DNA nanostructures. The LEGO gradient mixer presented here is an effective complement to the current state-of-the-art RZC method to purify DNA origami. The LEGO gradient mixer was able to produce a concentration gradient in a relatively short timespan compared to other gradient mixers.^27,30^ The gradient produced by this method was able to purify both 6-hb monomer from its staple strands and 6-hb dimer from its precursor monomer. Although the purified dimer shows only 63±6% purity (Fig. 5J), the observed monomer in the purified dimer comes from the difficulty of extracting the dimer sample manually from the small gradient volume in which the experiment is conducted. Taking note that we use less than ideal equipment for fractionation, *i.e.* long neck gel loading tips, the yield could significantly be improved by upgrading the fractionation technique. To achieve a consistent result, a better technique for fraction collection should be employed. It is also favorable to automate the fraction collection process using the currently available liquid handling equipment for critical applications. RZC purification method paired with the gradient mixing method presented in this paper could help pave the way for large-scale DNA nanostructure production that will enable separation of functional DNA nanostructures from the side products of bioconjugation reactions, such as excess of streptavidins, quantum dots, gold nanoparticles, aptamers, and other moeties.

We note that the reported method can be further improved to reduce the cost even further. The current cost of the EV3 LEGO Mindstorms pieces used to make this device is $349.99, while the currently available gradient mixer is quoted at around $500.00 to $650.00 (Millipore Sigma gradient mixer, cat no. Z340391) and >$2K for GradientMaster (Biocomp Instruments). However, the main components of the LEGO gradient mixer are essentially a motor, a servo, and a custom holder for the tubes, all of which do not need high precision. Hence, the price to create such a machine can be reduced further by an order of magnitude (Table S2). To lower the cost further, such machine can be powered manually using some gear and scaffolding materials. In the spirit of frugal science, our discovery could help density gradient centrifugation become a more accessible tool for scientific communities around the world. The myriad applications of density gradients to fractionate DNA origami, viral particles, and a variety of macromolecules^36^ have been critical in the preparation of viruses for vaccines and other immunotherapy products. The methods introduced in this paper will contribute to science education and research in resource-limited settings.

## Acknowledgements

The work was initiated by a class project in PHY 472 (Advanced Biophysics Laboratory) in the Department of Physics at Arizona State University. We acknowledge other students in PHY 472, Carter Swanson, Adrian Kwiatkowski, and Biruni Hariadi for valuable discussions. We thank Sapto Cahyono for assistance with illustrations, Evangeline Taylor-Hermes for pointing the LEGO software for the building instructions, and Gde Bimananda Wisna Mahardika for technical assistance. AFM images were collected in the lab of Hao Yan at Arizona State University. The authors also thank Jeffrey La Belle’s group for lending us the rotor used for ultra-centrifugation. The research in Hariadi lab was supported by the National Institutes of Health Director’s New Innovator Award (1DP2AI144247), National Science Foundation SemiSynBio II (2027215), and Arizona Biomedical Research Consortium (ADHS17-00007401).

## Supporting Information

## S1. Supplementary Figures

**Fig. S1.**
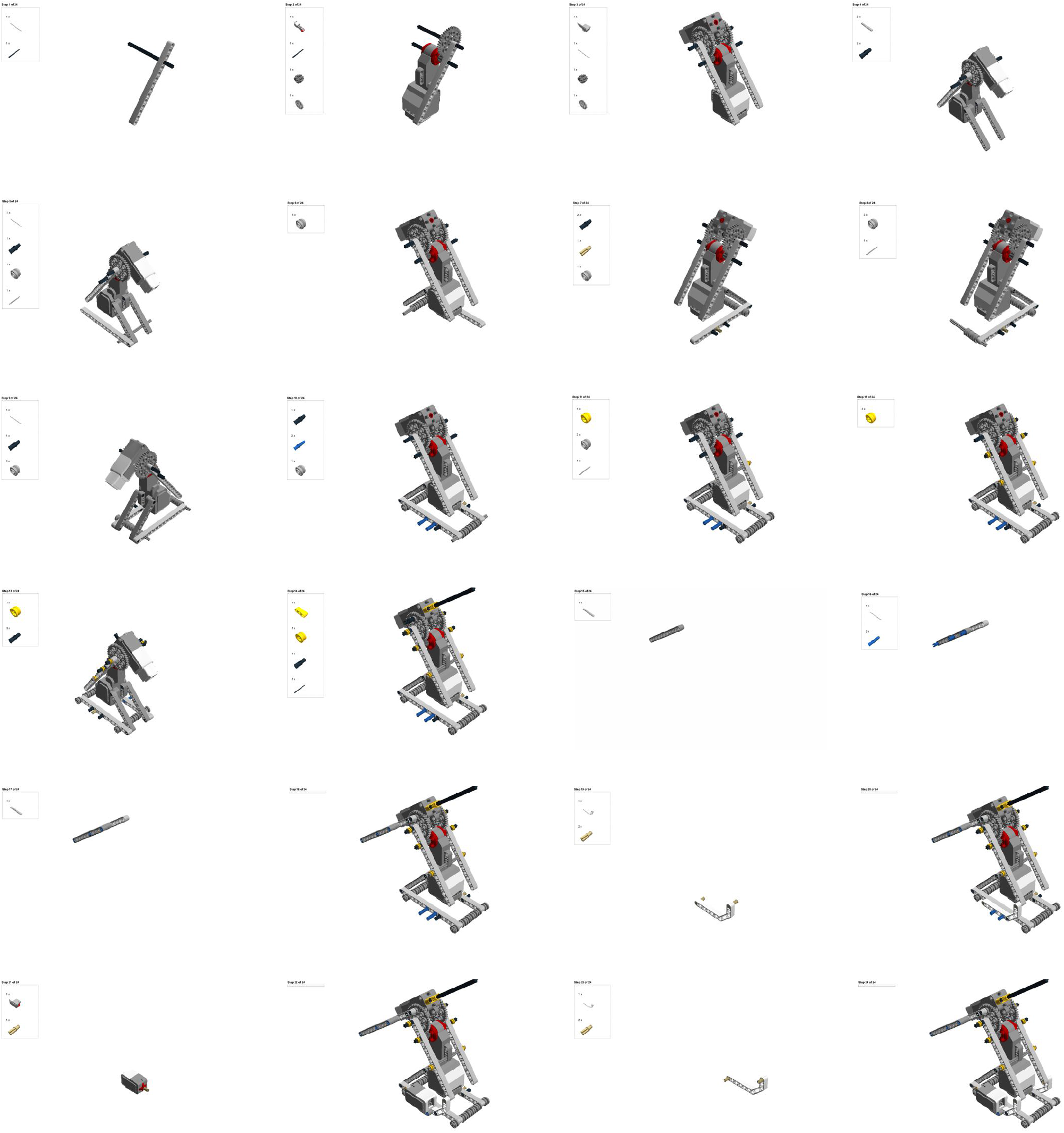
Building instruction for the LEGO gradient mixer (24 steps total) excluding the centrifuge tube holder (3D printed). Parts can be purchased from the EV3 LEGO Mindstorms kit.

**Fig. S2.**
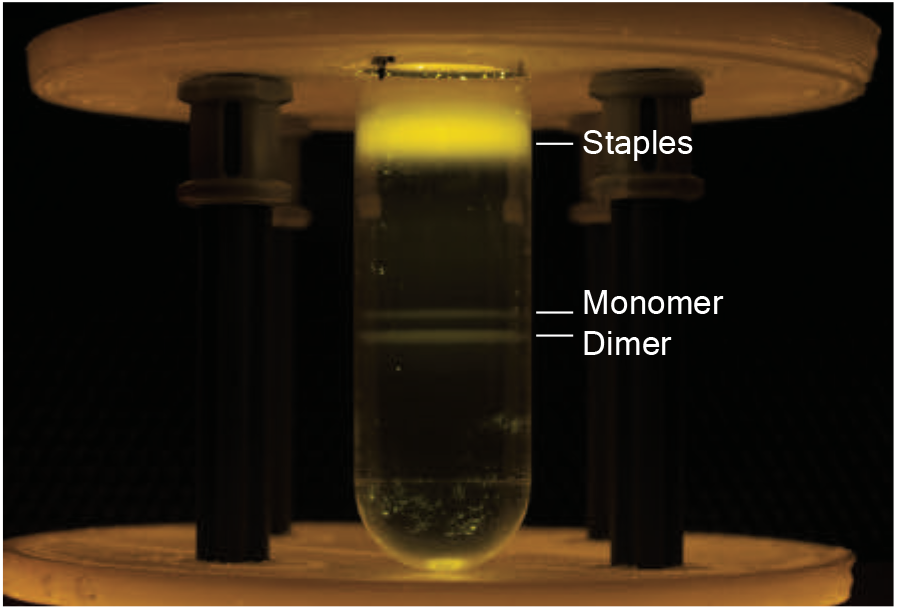
Separation of dimer by RZC using 30−60% (w/v) sucrose gradient created using LEGO gradient mixer. The sample was centrifuged at 50,000 rpm at 4°C for 6 hours.

**Fig. S3.**
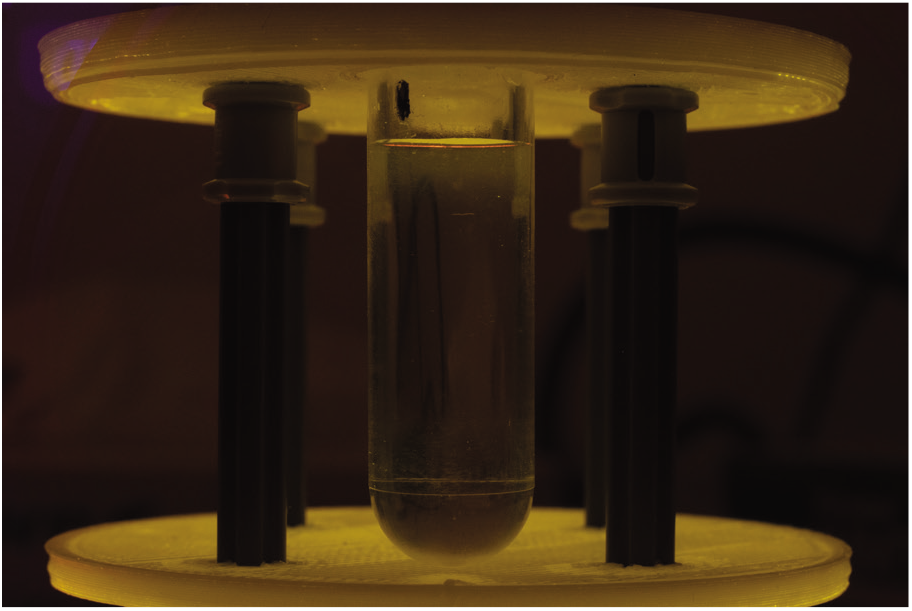
RZC of SYBR gold-only sample using 30−60% (v/v) glycerol gradient. The sample was centrifuged at 50,000 rpm at 4°C for 3 hours. The SYBR gold was undetectable under blue LED illumination.

## S2. DNA sequences

**Table S1.**
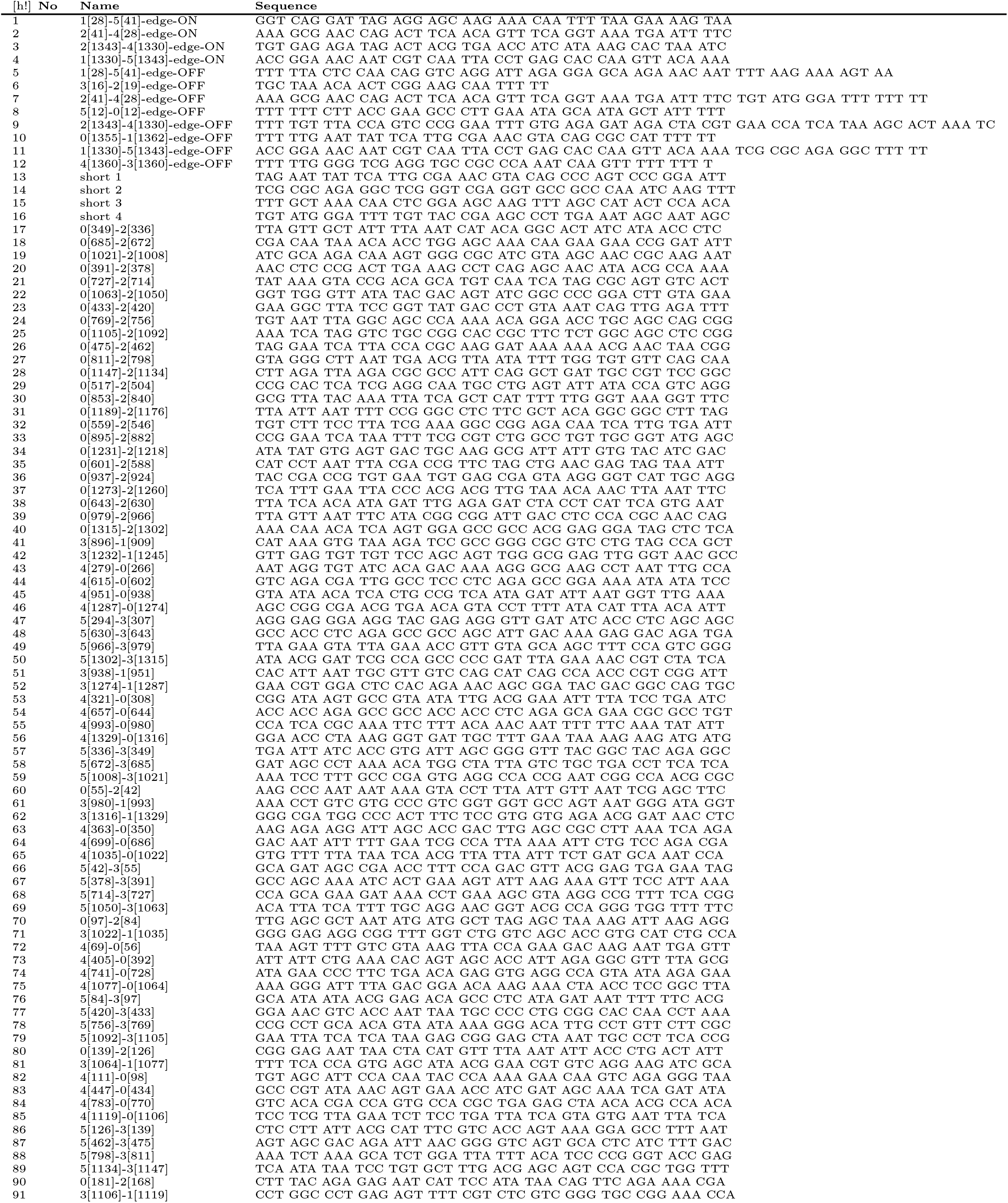
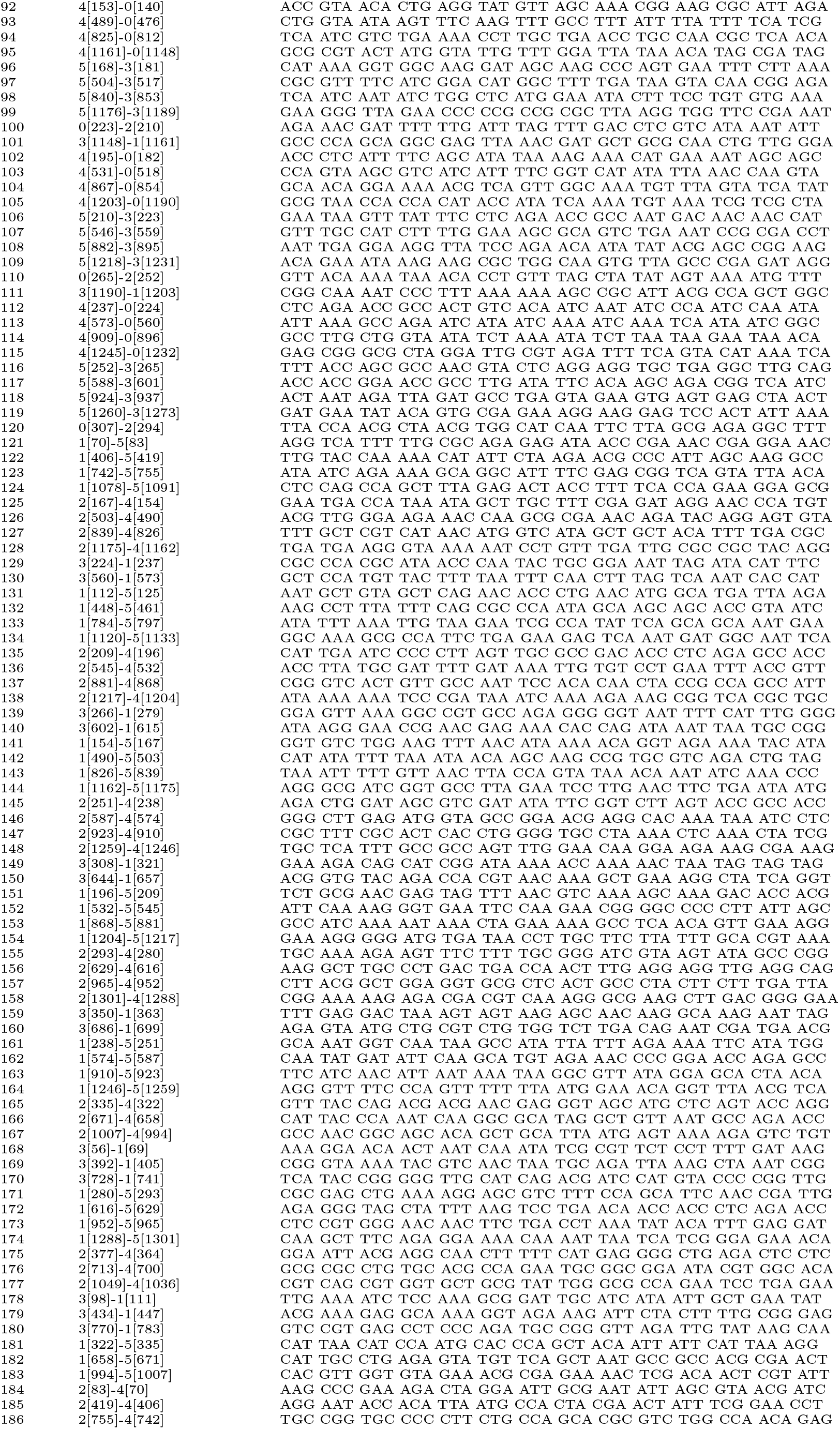
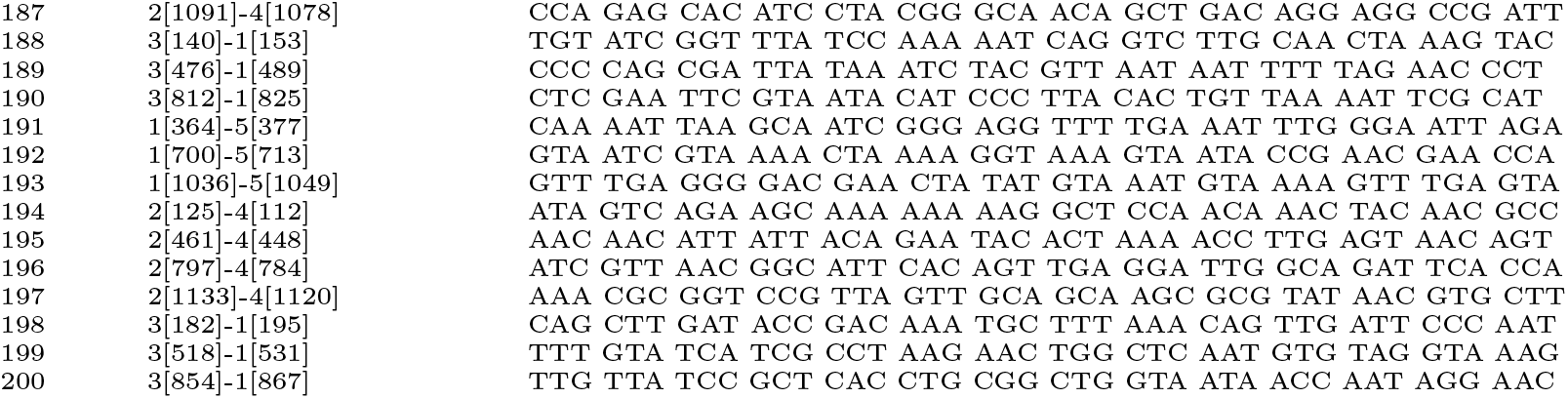
Computer aided staple strand sequences for the DNA origami nanotube monomers.

## S3. Cost analysis

**Table S2.** Estimated bill of materials for gradient mixer.

| Description | Purpose | Quantity | Price @ | Total price |
| --- | --- | --- | --- | --- |
| ELEGOO UNO R3 Board ATmega328P | Microcontroller | 1 | \$ 13.99 | \$ 13.99 |
| 6 VDC Mini motor | Motor | 1 | \$ 0.75 | \$ 0.75 |
| SG92R Mini servo | Servo | 1 | \$ 4.75 | \$ 4.75 |
| Qunqi L298N Motor Drive Controller Board | Motor Controller | 1 | \$ 6.69 | \$ 6.69 |
| OVERTURE PLA Filament | 3D Print Material | 300 gr | \$ 18.99 /kg | \$ 6.00 |
| **Total Price** | | | | **\$ 32.18** |

## S4. Protocols

### Reagents

**Concentration gradient**

- 50 × TAE buffer, pH 8.3 (VWR Life Science, cat. no. 75800-940)
- Magnesium chloride hexahydrate (Sigma-Aldrich, cat. no. M9272)
- Glycerol (VWR Life Science, cat. no. BDH1172)
- Sucrose (Sigma-Aldrich, product no. S 0389)

**DNA origami folding**

- 50× TAE buffer, pH 8.3 (VWR Life Science, cat. no. 75800-940)
- Magnesium chloride hexahydrate (Sigma-Aldrich, cat. no. M9272)
- Staple strands (Integrated DNA Technique Inc.)
- ssDNA scaffold p8064 (Tilibit Nanosystem, product no. M1-51)

**Agarose Gel Electrophoresis**

- SYBR gold DNA gel stain (Thermo Fisher Scientific, cat no. S11494)
- DNA gel loading dye (Thermo Fisher Scientific, cat no. R0631)
- 1 kb DNA ladder (Gold Biotechnology Inc., cat. no. D010-500)
- Agarose tablets (EURx, cat. no. E0305-01)

### Equipments

**General**

- P2L Eppendrof pipette (Gilson, SKU FA10001M)
- P10L Eppendrof pipette (Gilson, SKU FA10002M)
- P200L Eppendrof pipette (Gilson, SKU FA10005M)
- P1000L Eppendrof pipette (Gilson, SKU FA10006M)
- EXPERT tips E200 Tipack (Gilson, SKU F1733002)
- EXPERT Tips E1000 XL Tipack (Gilson, SKU F1735002)
- Milli-Q® Direct Water Purification System (Milipore Sigma, cat. no. ZR0Q016WW)

**Density gradient preparation**

- EV3 LEGO Mindstorm (LEGO, item no. 31313)
- Ultimaker 3 3D printer (part no. 9671)
- PLA 3D printing materials
- 1 mL syringe (Becton Dickinson, cat. no. 309628)

**DNA origami preparation**

- Thermocycler, MiniAmp™ Thermal Cycler REX (Thermo Fisher Scientific, cat No. A38080)
- PCR tubes (USA Scientific, cat. no. 1402-8120)
- 100 kD amicon filter (Milipore, cat. no. UFC510024)
- Benchmark MC-12 High Speed Microcentrifuge (Marshall Scientific, item no. C1612)

**Rate-zonal centrifugation**

- Beckman TLS 55 swinging bucket rotor (Beckman Coulter, part no. 346936)
- Polycarbonate centrifuge tube (Beckman Coulter, part no. 343778)
- Optima TLX ultracentrifuge (Beckman Coulter, part no. 361545)
- Longneck gel-loading tips (Fisher Scientific, cat. no. 02-707-81)
- Safe Imager 2.0 Blue Light transilluminators (Invitrogen, cat. no. G6600)

**Gel electrophoresis**

- 250 mL Erlenmeyer flask (Karter Scientific, part no. 214U2)
- External gel cast (Thermo Fisher Scientific, cat. no. B2-CST)
- Owl™ easyCast™ B2 Mini Gel Electrophoresis Systems (Thermo Fisher Scientific, cat. no. B2-BP)
- Bio-Rad Molecular Imager Gel Doc XR+ (Bio-Rad, part. no. 1708195EDU)

**Atomic force microscopy**

- MultiMode 8-HR AFM (Bruker)
- ScanAsyst-Fluid+probes cantilever (Bruker)
- Glass probe holder MTFML-V2 (Bruker)
- V1 AFM mica discs, 10mm (Ted Pella Inc., product no. 50-10)
- AFM/STM Metal Specimen Discs, 12mm (Ted Pella Inc., product no. 16208)

## Protocols

### DNA origami preparation

1 │ While referring to the staple strand sequences (Table S1), pool the staple strands into the following categories.

2 │ To create the 6-hb monomer mixture, combine 30 nm of the ssDNA p8064 scaffold strands and 300 nm each of the staple strands from Left OFF, Right OFF, and Core 6-hb. The strand mix should be added in 1× TAE 12.5 mM MgCl_2_ (final concentration).
3 │ To create the 6-hb dimer mixture, two types of precursor monomers mixture, *left* and *right*, need to be mixed in separate tubes. The *left* precursor monomers have complementary sequences with the *right* precursor monomer, which enable them to form a dimer. The *left* precursor monomer contains 30 nm of the ssDNA p8064 scaffold strands and 300 nm each of the staple strands in Left ON, Right OFF, Short 1 2, and Core 6-hb. The *right* precursor monomer contains 0 nm of the ssDNA p8064 scaffold strands and 300 nm each of the staple strands in Right ON, Left OFF, Short 3 4, and Core 6-hb. All origami mixture is in 1× TAE 12.5 mM MgCl_2_ (final concentration).
4 │ Anneal the 6-hb monomer, *left* precursor monomer, and *right* precursor monomer in separate PCR tubes using a thermocycler with a thermal gradient that starts at 90°C, gradually cools to 30°C over 1.5 hours, and ends with incubation at 4°C. <u>The 6-hb monomer is formed after this annealing process.</u> The 6-hb dimer needs an additional step after the *left* and *right* precursor monomers are formed.
5 │ Filter excess staple from the *left* and *right* monomer separately using 100 kD Amicon filtration on a centrifuge. Both precursor monomers should be filtered at 4,500 g for 5 minutes twice to ensure optimal staple filtration while maintaining the DNA origami structure integrity. After filtering the mixture twice, collect the DNA origami by flipping the amicon filter on a new tube and centrifuging them at 1,000 g for 2 minutes. **CRITICAL STEP** Filtration of excess staples from the precursor monomers is important to ensure the leftover staple from one precursor does not bind to the activated site of the other precursor, preventing dimerization by binding to the activated site.
6 │ Combine the *left* and *right* precursor monomer after filtration of excess staples in step 5. The combined mixture is stored at 4°C to dimerize overnight. <u>The 6-hb dimer is formed after this process.</u>
7 │ Once the 6-hb monomers and dimers are formed, perform an AGE analysis to check the formation of both origami. The monomer shows one band while the dimer shows two bands. The two bands in the dimer lane show one band at the same level as the monomer (from the precursor monomer that fails to dimerize) and another band slightly below the monomer (indicating the dimer).

| Category | Staple Strands |
| --- | --- |
| Left ON | staple 1–2 |
| Right ON | staple 3–4 |
| Left OFF | staple 5–8 |
| Right OFF | staple 9–12 |
| Short 1 2 | staple 13–14 |
| Short 3 4 | staple 15–16 |
| Core 6-hb | the rest of the staples |

### Preparation of density gradient

8 │ Assemble the LEGO gradient machine following the instructions in Figure S1. The LEGO parts needed to build the LEGO gradient mixer can be found in the EV3 LEGO Mindstorm pack.
9 │ 3D print the centrifuge tube holder file with PLA materials. The software used to produce the .gcode needed for 3D printing is Ultimaker Cura and the 3D printer used to print the centrifuge tube holder is Ultimaker 3. Any other 3D printer can also be used to print the centrifuge tube holder. Once the centrifuge tube holder is printed, attach the holder to the spinning motor (Figure 1b) of the LEGO assembly. The full assembly of the LEGO gradient mixer is shown in Figure 1.
10 │ Prepare several different concentrations of glycerol (or sucrose, depending on the viscous agent used). A common gradient used in the experiment is a 15%–45% (v/v) gradient.
11 │ Fill the centrifuge tube with 300 *μ*L of the 45% glycerol using a pipette. Make sure that the 45% glycerol is resting on the bottom of the tube and that no bubble is present. After the 45% glycerol, lay 300 *μ*L of 15% glycerol on top of the 45% glycerol in the centrifuge tube using a syringe to ensure minimal surface disturbance on the 45% glycerol layer. The border between the 15% and 45% glycerol should be visible.
12 │ Load the glycerol-filled centrifuge tube into the designated holder in the LEGO gradient mixer. Run the LEGO_gradient_mixer_protocol.ev3 protocol found in the Supplementary Materials Repository, EV3-protocols folder. The protocol will tilt the glycerol filled tube 90°to a horizontal position and spin the tubes for one minute to mix the two different glycerol concentrations. After spinning the tubes, the LEGO gradient mixer will return the tubes to its initial position. A faded boundary and smooth gradient between the 15% and 45% glycerol can now be observed.

### Rate Zonal Centrifugation and Fractionation

13 │ Prepare the sample for RZC by mixing it with glycerol to a 10% (v/v) final concentration. Adding glycerol to the sample ensures the sample sits on the top surface of the glycerol gradient and eases the sample’s entry into the glycerol gradient during RZC.
14 │ Carefully load 50-100 *μ*L of the sample (10% glycerol) on top of the gradient using a pipette, making sure the sample does not pierce the gradient. Most of the sample should rest on the surface of the gradient.
15 │ Insert the centrifuge tubes containing the sample and gradient into a swinging-bucket rotor (Beckman TLS 55). **CRITICAL STEP** A swinging-bucket rotor should be used instead of a fixed-angle or vertical rotor because the glycerol gradient used is positioned from top to bottom. To prevent the sample from pelleting on the side and to ensure that the sample travels down the gradient, a swinging bucket rotor is needed.
16 │ Centrifuge the rotor at 50,000 rpm (~150,000 g) for 1.5 hours at 4°C in the Optima TLX ultracentrifuge. For different origami, the RPM and duration of centrifugation need to be optimized.
17 │ Once the centrifugation is complete, collect 16 to 18 equal volume fractions of the gradient containing the sample. Longneck gel loading tips can be used with a pipette to fractionate the gradient from bottom to top. For a manual method such as this, fractionation from bottom to top is recommended because it ensures a more organized way to control the position of the tips. Other methods of fractionation using liquid handling instruments are preferred to achieve a more consistent RZC result.

### Agarose Gel Electrophoresis

18 │ After the sample has been fractionated, take 10 μL from each fraction and mix it with 1 μL of 20× SYBR gold and 1 μL of loading dye. Leave it to incubate at room temperature for 30 min.
19 │ Cast a 1% agarose gel with enough lanes to accommodate the number of fractions being analyzed.
20 │ When the gel is ready to use, put the gel into its electrophoresis chamber and fill the chamber with 1× TAE 12.5 mM MgCl_2_ buffer until the gel is submerged.
21 │ Load 1 kb DNA ladder and all fractions that have been labelled with SYBR gold (step 18) into their respective lanes in the gel. Start the electrophoresis using a 75 V power supply for 1.5 hours.
22 │ After the gel electrophoresis is done, image the gel using Bio-Rad Molecular Imager Gel Doc XR+. Set the setting to image nucleic acids with SYBR Gold label. The bands in the gel show the staple strands near the top fraction lanes and the origami somewhere in the middle to bottom fractions. If the bands are not immediately clear, post-staining the gel in 1× SYBR Gold for 15–30 minutes will increase the band intensity.
23 │ Combine the fractions containing the origami (based on the AGE result) into a single tube which consists of the purified origami.

### AFM imaging

24 │ Turn on the MultiMode 8 controller and AFM to warm up the laser.
25 │ Mount the 10 mm V1 mica discs onto the AFM/STM 12 mm metal specimen discs and cleave the mica sheet with a tape to an atomically-flat surface for AFM imaging.
26 │ Dilute the fractionated sample by 100× with 1× TAE 12.5 mM MgCl_2_. Depending on the purified sample concentration, the dilution factor may need to be adjusted.
27 │ Pipette 30 μL of the diluted sample onto the freshly cleaved mica. More sample is sometimes added to cover the mica surface. Place the mica puck containing the sample inside the AFM.
28 │ Mount the scanAsyst-Fluid+probes cantilever to the glass probe holder MTFML-V2 with a tweezer and insert them into the AFM.
29 │ Lower the cantilever until it touches the sample and adjust the laser position and mirror to reach a gain of at least 6.00.
30 │ Calibrate the zero position for the vertical and horizontal position and engage the cantilever to begin scanning. The scanned images are flattened and the length of the 6-hb origami(s) is measured to determine the presence of monomers (~500 nm) and dimers (~1000 nm).

## S5. Supplementary Materials Repository

LEGO assembly instructions and all data supporting the findings of this study are available within the paper and in https://github.com/jsentosa/LEGO-gradient-maker.git. All raw data and AFM images are available upon request.

**Movie S1** Gradient formation using the LEGO robot. https://github.com/jsentosa/LEGO-gradient-maker/tree/master/Recording-protocols

## References

1. Paul W K Rothemund. Folding DNA to create nanoscale shapes and patterns. Nature, 440(7082):297, 2006.

2. Ebbe S Andersen, Mingdong Dong, Morten M Nielsen, Kasper Jahn, Ramesh Subramani, Wael Mamdouh, Monika M Golas, Bjoern Sander, Holger Stark, Cristiano LP Oliveira, Jan Skov Pedersen, Victoria Birkedal, Flemming Besenbacher, Kurt V Gothelf, and Jørgen Kjems. Self-assembly of a nanoscale DNA box with a controllable lid. Nature, 459(7243):73–76, 2009.

3. Shawn M Douglas, Hendrik Dietz, Tim Liedl, Björn Högberg, Franziska Graf, and William M Shih. Self-assembly of DNA into nanoscale three-dimensional shapes. Nature, 459(7245):414–418, 2009.

4. Hendrik Dietz, Shawn M Douglas, and William M Shih. Folding DNA into twisted and curved nanoscale shapes. Science, 325(5941):725–730, 2009.

5. Dongran Han, Suchetan Pal, Yan Liu, and Hao Yan. Folding and cutting DNA into reconfigurable topological nanostructures. Nat Nanotechnol, 5(10):712–717, 2010.

6. Tim Liedl, Björn Högberg, Jessica Tytell, Donald E Ingber, and William M Shih. Self-assembly of three-dimensional prestressed tensegrity structures from DNA. Nat Nanotechnol, 5(7):520–524, 2010.

7. Dongran Han, Suchetan Pal, Jeanette Nangreave, Zhengtao Deng, Yan Liu, and Hao Yan. DNA origami with complex curvatures in three-dimensional space. Science, 332(6027):342–346, 2011.

8. Mark Bathe and Paul W K Rothemund. DNA nanotechnology: A foundation for programmable nanoscale materials. MRS Bull, 42(12):882–888, 2017.

9. Jeanette Nangreave, Dongran Han, Yan Liu, and Hao Yan. DNA origami: a history and current perspective. Curr Opin Chem Boil, 14(5):608–615, 2010.

10. William M Shih and Chenxiang Lin. Knitting complex weaves with DNA origami. Curr Opin Struc Biol, 20(3):276–282, 2010.

11. Thomas Tørring, Niels V Voigt, Jeanette Nangreave, Hao Yan, and Kurt V Gothelf. DNA origami: a quantum leap for self-assembly of complex structures. Chem Soc Rev, 40(12):5636–5646, 2011.

12. Rémi Veneziano, Tyson J Moyer, Matthew B Stone, Eike-Christian Wamhoff, Benjamin J Read, Sayak Mukherjee, Tyson R Shepherd, Jayajit Das, William R Schief, Darrell J Irvine, and Mark Bathe. Role of nanoscale antigen organization on b-cell activation probed using DNA origami. Nat Nanotechnol, 15(8):716–723, 2020.

13. Qian Zhang, Qiao Jiang, Na Li, Luru Dai, Qing Liu, Linlin Song, Jinye Wang, Yaqian Li, Jie Tian, Baoquan Ding, and Yang Du. DNA origami as an in vivo drug delivery vehicle for cancer therapy. ACS Nano, 8(7):6633–6643, 2014.

14. Qiao Jiang, Chen Song, Jeanette Nangreave, Xiaowei Liu, Lin Lin, Dengli Qiu, Zhen-Gang Wang, Guozhang Zou, Xingjie Liang, Hao Yan, and Baoquan Ding. DNA origami as a carrier for circumvention of drug resistance. J Am Chem Soc, 134(32):13396–13403, 2012.

15. Yong-Xing Zhao, Alan Shaw, Xianghui Zeng, Erik Benson, Andreas M Nyström, and Björn Högberg. DNA origami delivery system for cancer therapy with tunable release properties. ACS Nano, 6(10):8684–8691, 2012.

16. Joona Mikkila, Antti-Pekka Eskelinen, Elina H Niemela, Veikko Linko, Mikko J Frilander, Päivi Törmä, and Mauri A Kostiainen. Virus-encapsulated DNA origami nanostructures for cellular delivery. Nano Lett, 14(4):2196–2200, 2014.

17. Christian Steinhauer, Ralf Jungmann, Thomas L Sobey, Friedrich C Simmel, and Philip Tinnefeld. DNA origami as a nanoscopic ruler for super-resolution microscopy. Angew Chem Int Edit, 48(47):8870–8873, 2009.

18. Jürgen J Schmied, Mario Raab, Carsten Forthmann, Enrico Pibiri, Bettina Wünsch, Thorben Dammeyer, and Philip Tinnefeld. DNA origami–based standards for quantitative fluorescence microscopy. Nat Protoc, 9(6):1367–1391, 2014.

19. Sarah Helmig, Alexandru Rotaru, Dumitru Arian, Larisa Kovbasyuk, Jacob Arnbjerg, Peter R Ogilby, Jørgen Kjems, Andriy Mokhir, Flemming Besenbacher, and Kurt V Gothelf. Single molecule atomic force microscopy studies of photosensitized singlet oxygen behavior on a DNA origami template. ACS Nano, 4(12):7475–7480, 2010.

20. Ralf Jungmann, Christian Steinhauer, Max Scheible, Anton Kuzyk, Philip Tinnefeld, and Friedrich C Simmel. Single-molecule kinetics and super-resolution microscopy by fluorescence imaging of transient binding on DNA origami. Nano Lett, 10(11):4756–4761, 2010.

21. Nadrian C Seeman. DNA engineering and its application to nanotechnology. Trends Biotechnol, 17(11):437–443, 1999.

22. Albert M Hung, Christine M Micheel, Luisa D Bozano, Lucas W Osterbur, Greg M Wallraff, and Jennifer N Cha. Large-area spatially ordered arrays of gold nanoparticles directed by lithographically confined DNA origami. Nat Nanotechnol, 5(2):121–126, 2010.

23. Veikko Linko, Seppo-Tapio Paasonen, Anton Kuzyk, Päivi Törmä, and J Jussi Toppari. Characterization of the conductance mechanisms of DNA origami by ac impedance spectroscopy. Small, 5(21):2382–2386, 2009.

24. Yonggang Ke, Gaëtan Bellot, Niels V Voigt, Elena Fradkov, and William M Shih. Two design strategies for enhancement of multilayer–DNA-origami folding: underwinding for specific intercalator rescue and staple-break positioning. Chemical Sci, 3(8):2587–2597, 2012.

25. Klaus F Wagenbauer, Floris AS Engelhardt, Evi Stahl, Vera K Hechtl, Pierre Stömmer, Fabian Seebacher, Letizia Meregalli, Philip Ketterer, Thomas Gerling, and Hendrik Dietz. How we make DNA origami. ChemBioChem, 18(19):1873–1885, 2017.

26. Alan Shaw, Erik Benson, and Björn Högberg. Purification of functionalized DNA origami nanostructures. ACS Nano, 9(5):4968–4975, 2015.

27. Chenxiang Lin, Steven D Perrault, Minseok Kwak, Franziska Graf, and William M Shih. Purification of DNA-origami nanostructures by rate-zonal centrifugation. Nucleic Acids Res, 41(2):e40, 2012.

28. Christopher Timm and Christof M Niemeyer. Assembly and purification of enzyme-functionalized DNA origami structures. Angew Chem Int Edit, 54(23):6745–6750, 2015.

29. R J Britten and R B Roberts. High-resolution density gradient sedimentation analysis. Science, 131(3392):32–33, 1960.

30. Uwe Michelsen and Jörg Von Hagen. Chapter 19 isolation of subcellular organelles and structures. Method Enzymol, page 305–328, 2009.

31. Shawn M Douglas, James J Chou, and William M Shih. DNA-nanotube-induced alignment of membrane proteins for NMR structure determination. P Natl Acad Sci USA, 104(16):6644–6648, 2007.

32. Jonathan F Berengut, Julian C Berengut, Jonathan P K Doye, Domen Prešern, Akihiro Kawamoto, Juanfang Ruan, Madeleine J Wainwright, and Lawrence K Lee. Design and synthesis of pleated DNA origami nanotubes with adjustable diameters. Nucleic Acids Res, 47(22):11963–11975, 2019.

33. Florian Praetorius, Benjamin Kick, Karl L Behler, Maximilian N Honemann, Dirk Weuster-Botz, and Hendrik Dietz. Biotechnological mass production of DNA origami. Nature, 552(7683):84–87, 2017.

34. Stefan Niekamp, Katy Blumer, Parsa M Nafisi, Kathy Tsui, John Garbutt, and Shawn M Douglas. Folding complex DNA nanostructures from limited sets of reusable sequences. Nucleic Acids Res, 44(11):e102, 2016.

35. Jean-Philippe J Sobczak, Thomas G Martin, Thomas Gerling, and Hendrik Dietz. Rapid folding of DNA into nanoscale shapes at constant temperature. Science, 338(6113):1458–1461, 2012.

36. Mohamed A Desai and Sandra P Merino. Application of density gradient ultracentrifugation using zonal rotors in the large-scale purification of biomolecules. In Downstream Processing of Proteins, pages 59–72. Springer, 2000.

